# Molecular basis of selective cytokine signaling inhibition by antibodies targeting a shared receptor

**DOI:** 10.1101/2021.05.07.443154

**Authors:** James K. Fields, Kyle Kihn, Gabriel S. Birkedal, Erik H. Klontz, Kjell Sjöström, Sebastian Günther, Robert Beadenkopf, Göran Forsberg, David Liberg, Greg A. Snyder, Daniel Deredge, Eric J. Sundberg

## Abstract

Interleukin-1 (IL-1) family cytokines are potent mediators of inflammation, acting to coordinate local and systemic immune responses to a wide range of stimuli. Aberrant signaling by IL-1 family cytokine members, however, is linked to myriad inflammatory syndromes, autoimmune conditions and cancers. As such, blocking the inflammatory signals inherent to IL-1 family signaling is an established and expanding therapeutic strategy. While several FDA-approved IL-1 inhibitors exists, including an Fc fusion protein, a neutralizing antibody, and an antagonist cytokine, none specifically targets the co-receptor IL-1 receptor accessory protein (IL-1RAcP). Most IL-1 family cytokines form productive signaling complexes by binding first to their cognate receptors – IL-1RI for IL-1α and IL-1β; ST2 for IL-33; and IL-36R for IL-36α, IL-36β and IL-36γ – after which they recruit the shared secondary receptor IL-1RAcP to form a ternary cytokine/receptor/co-receptor complex. Recently, IL-1RAcP was identified as a biomarker for both AML and CML. IL-1RAcP has also been implicated in tumor progression in solid tumors and an anti-IL1RAP antibody (nadunolimab, CAN04) is in phase II clinical studies in pancreatic cancer and non-small cell lung cancer (NCT03267316). As IL-1RAcP is common to all of the abovementioned IL-1 family cytokines, targeting this co-receptor raises the possibility of selective signaling inhibition for different IL-1 family cytokines. Indeed, previous studies of IL-1β and IL-33 signaling complexes have revealed that these cytokines employ distinct mechanisms of IL-1RAcP recruitment even though their overall cytokine/receptor/co-receptor complexes are structurally similar. Here, using functional, biophysical, and structural analyses, we show that antibodies specific for IL-1RAcP can differentially block signaling by IL-1 family cytokines depending on the distinct IL-1RAcP epitopes that they engage. Our results indicate that targeting a shared cytokine receptor is a viable therapeutic strategy for selective cytokine signaling inhibition.

## Introduction

Interleukin-1 (IL-1), a critical regulator of inflammation and pro-inflammatory cytokine of the IL-1 superfamily, has long been known to play a role in inflammatory syndromes, autoimmune diseases, and cancers [1-6]. Due to the vast number of organs and cells that IL-1 affects, from the thalamus to T lymphocytes, it is unsurprising that the list of IL-1-mediated syndromes is long, including joint and muscle diseases, such as rheumatoid arthritis (RA), osteoarthritis, and osteomyelitis; hereditary autoinflammatory conditions such as familial Mediterranean fever (FMF) and cryopyrin associated periodic syndrome (CAPS); systemic inflammatory diseases including macrophage activation syndrome and Still’s disease; as well as common diseases including Gout, type 2 diabetes, Hidradenitis suppurativa, and cancer. [7, 8]. Importantly, it has been shown that blocking IL-1 signaling leads to a reduction and/or reversal of many IL-1-mediated diseases [9-14].

Inflammatory IL-1 signaling is a stepwise process regulated at multiple levels, from gene expression to the inhibitory actions of antagonist cytokines and decoy receptors [3]. First, IL-1 binds with high affinity to its primary receptor interleukin-1 receptor 1 (IL-1RI). Next, the cytokine/receptor complex (IL-1/IL-1RI) recruits the common co-receptor Interleukin-1 receptor accessory protein (IL-1RAcP). As this ternary complex (IL-1/IL-1RI/IL-1RAcP) forms, Toll/Interleukin-1 receptor (TIR) domains, attached through single transmembrane helices to each receptor, engage one another intracellularly, initiating a potent signaling cascade. Blocking this inflammatory signal at the cytokine/receptor level is currently the most common and effective therapeutic strategy.

So far, only three IL-1 targeting therapeutics are licensed for widespread use: Anakinra, the natural IL-1 antagonist; Rilonacept, an Fc-fused decoy receptor; and Canakinumab, an IL-1β neutralizing antibody. While Anakinra is the gold-standard for treating IL-1 mediated diseases, its 4-to 6-hour therapeutic half-life and daily injection schedule makes it less than ideal for long term treatment. Rilonacept must also be administered on a weekly schedule. Indeed, there is pressing need for additional IL-1 therapeutics with longer half-lives, such as canonical IgG molecules. The efficacy of Canakinumab, one such molecule, in a multitude of syndromes is supportive of this idea. To date, a receptor targeting antibody to block IL-1 signaling is not in wide use for IL-1-mediated diseases.

Within the IL-1 family of cytokines, of which there are seven agonist cytokines, both IL-1 and IL-33 share a common co-receptor: IL-1RAcP. While IL-1RAcP is primarily implicated in innate and adaptive immunity, this receptor was reported as a biomarker for chronic myeloid leukemia (CML) stem cells in 2010 [8]. Prior to this finding, no cell surface marker existed that permitted the separation of normal hematopoietic stem cells (HSCs) and CML cells [8]. The discovery of IL-1RAcP over-expression was extended to acute myeloid leukemia (AML) in 2012 [15]. Although the function of IL-1RAcP on AML and CML cells is still being investigated, IL-1 signaling has been shown to help AML and CML cells proliferate. Conversely, this proliferation is mitigated by the administration of the IL-1 receptor antagonist (IL-1Ra) [16-18].

Antibodies were subsequently raised against IL-1RAcP and tested for therapeutic benefit against AML and CML in xenograft mouse models [19, 20]. The IL-1RAcP-specific antibody CAN04, previously referred to as mAb3F8, exhibits strong antileukemic effect *in vivo*. CAN04 targets AML and CML cells for destruction through antibody-dependent cellular cytotoxicity (ADCC) [8, 21]. In addition, CAN04 is able to suppress AML and CML cell proliferation through the targeting of IL-1RAcP and subsequent blocking of IL-1 [19, 20]. CAN03, a second IL-1RAcP specific antibody, was developed as a backup candidate to CAN04.

Targeting a shared co-receptor in order to inhibit signaling, however, raises the question as to the antibody’s effect on signaling by the diverse members of the IL-1 cytokine family: If an antibody targets a shared IL-1 family signaling co-receptor, will it inhibit just IL-1? Or will it also affect IL-33 signaling? And, what is the feasibility of targeting a shared co-receptor to selectively inhibit signaling by cytokines that require the same co-receptor? Here, we show that the inhibitory potency and cytokine specificity of an anti-IL-1RAcP antibody depends predominantly on the location and composition of its epitope on the IL-1RAcP co-receptor. Using a range of functional, structural, and biophysical analyses, we provide the molecular basis for how two anti-IL-1RAcP antibodies, CAN03 and CAN04, exert distinct potencies in IL-1β inhibition while both preferentially inhibit IL-1β versus IL-33 signaling.

## Materials and Methods

### Protein Expression and Purification

All cytokines, antagonist cytokines, and soluble receptors were cloned, expressed, and purified as described in Günther *et al*. [22]. Briefly, proteins were purified by immobilized metal affinity chromatography (IMAC; HisPur NiNTA resin, ThermoFisher Scientific). The His-tag was removed by Tobacco Etch Virus (TEV) protease digestion and untagged cytokines were further purified by reverse NiNTA purification. IL-1RAcP was expressed by transiently transfecting HEK293T cells. After transfection using PEI (350ug PEI per 112ng DNA), cells were cultured for 84hr in FreeStyle F17 medium, supplemented with glutamax (1/100) and geneticin (1/1000). For the generation of a high-mannose variant of IL-1RAcP, Freestyle F17 medium was supplemented with kifunensine (1ug/mL). Proteins were purified from the supernatant using IMAC (HisTrap Excel, GE Healthcare). For deglycosylated variants, high-mannose IL-1RAcP glycans were cleaved using Endoglycosidase A (EndoA). Lastly, proteins were purified by size-exclusion chromatography (SEC) in 20mM HEPES, 150mM NaCl, pH 7.4 (cytokines using Superdex 75 and receptors using Superdex200, prep grade media, GE Healthcare). In addition, all cytokine purifications contained 2mM DTT. The antibodies CAN03 and CAN04 (murine IgG2a) were provided by Cantargia AB. Fabs were obtained by papain (ThermoFisher Scientific: cat20341) digestion of full length CAN03 and CAN04, followed by a subsequent reverse protein A purification and a Superdex 75 SEC column.

### Surface Plasmon Resonance

Affinities and kinetic parameters of protein-protein interactions were measured by surface plasmon resonance (SPR) analysis using a Biacore T100 biosensor (GE Healthcare). 2000 response units (RU) of Protein A from *Staphylococcus aureus* (Sigma Aldrich) were immobilized on all channels of a CM5 sensor chip. Approximately 200 RU of antibodies CAN03 and CAN04 were directly captured on flow cell 2. IL-1RAcP was then used as the analyte and titrated over flow cells 1 & 2 in two-fold dilutions. Sensorgrams were double-referenced against the control flow cell and buffer injections. Data were fit to a 1:1 binding model using Biacore T100 Evaluation Software.

### Cell-based signaling inhibition

To measure the potency of the antagonists and antibodies, HEK293T cells were transiently transfected with a luciferase gene (nano-luc, plasmid pNL2.2, Promega) under the control of the IL-8 promoter [23]. For measurement of IL-1β activity, only the reporter gene was transfected into the cells since IL-1RAcP and IL-1RI are endogenously expressed by HEK293T cells [24]. For measurement of IL-33 activity, both the reporter gene and the full-length ST2 gene were transfected (with a mass ratio of reporter to receptor plasmid of 25:1). 18hr after transfection, cells were harvested and seeded into 96-well plates at a concentration of 40,000 cells/well. Cells were stimulated with 5 pM IL-33 or 100 pM IL-1β. These concentrations were shown to activate IL-33 and IL-1 signaling to 90% of full activation. After 5 hr stimulation at 37C, cells were lysed and luciferase activity was determined using a Veritas luminescence reader (Promega). As a negative control, luciferase activity without cytokine stimulation was measured.

### Hydrogen-Deuterium Exchange Coupled to Mass Spectrometry (HDX-MS)

Peptide identification and coverage maps for IL-1RAcP, CAN03, and CAN04 were obtained from undeuterated controls as follows: 1 µL of 20 µM protein in 20 mM HEPES pH 7.4, 150 mM NaCl was diluted with 19 µL of ice cold quench (50 mM Glycine pH 2.4, 6.8 M Guanidine-HCl, 100mM TCEP) for 1 min prior to dilution with 180 µL of Buffer A (0.1% formic acid) and injection into a Waters HDX nanoAcquity UPLC (Waters, Milford, MA) with in-line protease XIII/pepsin digestion (NovaBioAssays). Peptic fragments were trapped on an Acquity UPLC BEH C18 peptide trap and separated on an Acquity UPLC BEH C18 column. A 7 min, 5% to 35% acetonitrile (0.1% formic acid) gradient was used to elute peptides directly into a Waters Synapt G2-Si mass spectrometer (Waters, Milford, MA). MS data were acquired with a 20 to 30 V ramp trap CE for high energy acquisition of product ions as well as continuous lock mass (Leu-Enk) for mass accuracy correction. Peptides were identified using the ProteinLynx Global Server 3.0.3 (PLGS) from Waters. Further filtering of 0.3 fragments per residues was applied in DynamX 3.0.

For each protein and complex, the HD exchange reactions were acquired using a LEAP autosampler controlled by Chronos software. The reactions were performed as follows: 2 µL of protein was incubated in 18 µL of 20 mM HEPES 99.99% D_2_O, pD 7.4, 150 mM NaCl. The following protein concentrations were used: 20 µM CAN03, 20 µM CAN04, 20 µM IL-1RAcp, 8.5 µM CAN03/IL-1RAcp complex and 8.5 µM CAN04/IL-1RacP complex. All reactions were performed at 25°C. Prior to injection, deuteration reactions were quenched at various times (10 sec, 1 min, 10 min, 1 hr and 2hr) with 60 µL of 50 mM Glycine buffer pH 2.4, 6.8 M Guanidine-HCl, 100 mM TCEP for 3 minutes prior to dilution with 200 µL of Buffer A. Subsequently, 250 µL of sample was injected and LC/MS acquisition performed as for the undeuterated controls. All deuteration time points were acquired in triplicate. Spectral curation, centroid calculation and deuterium uptake for all identified peptides with increasing deuterium incubation time were performed using Water’s DynamX 3.0 software.

### Molecular modeling of antibody-IL-1RAcP complexes

Docking poses were generated between the receptor IL-1RAcP (pdb: 4DEP) and the Fv regions of CAN03 and CAN04 (modeled using ABodyBuilder Fv prediction) [25]. Models of the docked structures, IL-1RAcP to CAN03 and IL1-RAcP to CAN04, were generated using the PatchDock web server [26]. For each docking simulation, the likely binding region for each receptor and ligand, as determined by the HDX-MS data, were added as a scoring parameter to PatchDock. The clustering RMSD was set to the default 4Å. For each simulation, the top 100 poses were extracted. For each docked system, poses were analyzed for CDR usage, IL-1RAcP binding, and alanine scan data. The top two poses for each docking pair were selected for HDX modeling. HDX modeling was performed using the ‘calc-HDX’ function of the HDXer tool with the given docked structure as input [27]. To calculate deuterium uptake, the Best & Vendruscolo phenomenological equation (Eq. 1) was used to calculate protection factors at individual backbone amide hydrogen through the course of a given simulation [28].

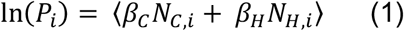

The protection factor (*P*_*i*_) at residue i is the ensemble average of the sum of the number of non-hydrogen atoms within 6.5 Åof the backbone Nitrogen atom of the residue (*N*_*C,i*_) multiplied by a scaling factor (*β*_*C*_) and the number of hydrogen bonds with the backbone amide hydrogen of the residues (*N*_*H,i*_) multiplied by a scaling factor (*β*_*H*_). In the calculation of (*N*_*C,i*_), the non-hydrogen atoms of the neighboring two residues on each side of the residue were omitted. Scaling factors of 0.35 and 2.0 were used for *β*_*C*_ and (*β*_*H*_) respectively. Protection factors were then used to calculate peptide level deuterium fractional uptake 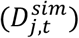 as a function of time point (t) of exchange (Eq. 2):

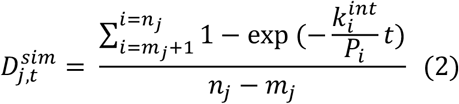

Where m_j_ and n_j_ are the starting and ending residue number of the i^th^ peptide and are chosen to match the experimental peptide segments observed in HDX-MS. Prolines, which do not have a backbone amide hydrogen, and the first residue of each segment were omitted from the calculations. The empirically determined intrinsic rate of exchange, k_int_ was then obtained [29]. In order to compare to the experimental data, the deuterium fractional uptake 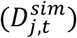 was transformed to deuterium uptake by multiplying 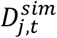 by the maximum theoretical number of exchangeable amide hydrogens. Difference plots for the modeled HDX were created for each pose by subtracting the modeled HDX exchange of the apo IL-1RAcP structure from the HDX exchange of the bound IL-1RAcP. RMSD values between the experimentally derived difference plots and the modeled different plots were then calculated (Eq. 3):

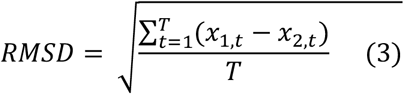

where t is an individual time point, x_1_ is the modeled data point, x_2_ is the experimental data point, and T is the total number of time points.

### ELISA Measurements in Chimeric IL-1RAcP Binding Studies

Microtiter plates were coated with 100 ng of mouse, human or chimeric mouse/human IL-1RAcP (100 µl/well) diluted in 0.01 M PBS, pH 7.4, and incubated overnight at 4°C. Plates were washed with ELISA washing buffer (0.01 M PBS, 0.05% Tween 20, pH 7.4) followed by a blocking step using 150 µl/well of ELISA blocking solution (PBS, 0.5% BSA, 0.05% Tween 20, pH 7.4). After one hour incubation at room temperature on agitation the plates were washed again using ELISA washing buffer. A series of CAN04 dilutions in ELISA blocking solution was prepared (ranging from 0.3 to 5000 ng/mL) and then transferred to the wells at 100 µl/well. The polyclonal antibodies KMT-2 (affinity purified rabbit polyclonal antibodies against hIL-1RAcP) and KMT-3 (affinity purified rabbit polyclonal antibodies against mIL-1RAcP) were diluted similarly and used as controls. Plates were incubated at room temperature for one hour on agitation and then washed with ELISA washing solution. One hundred µl/well of goat anti-human IgG conjugated to alkaline phosphatase was added and incubated one hour at room temperature on agitation. For the controls KMT-2 and KMT-3, goat anti-rabbit IgG conjugated to alkaline phosphatase was used. The plates were washed using ELISA washing solution followed by addition of substrate (4-Nitrophenyl phosphate disodium salt hexahydrate, Sigma-Aldrich, 1 mg/ml), 150 µl/well. The plates were thereafter incubated at room temperature on agitation and absorbance at 405 nm was measured after 30 min. Absorbance at 0 min was taken as background signal.

### Alanine Scanning mutagenesis

Alanine mutants were generated by site-directed mutagenesis (Qiagen) of Fc-fused IL-1RAcP and the procurement of the alanine library from Günther *et al*. [22]. Kinetic parameters and affinities of protein-protein interactions were measured by surface plasmon resonance (SPR) analysis using a Biacore T100 biosensor (GE Healthcare). 2000 response units (RU) of Protein A from *Staphylococcus aureus* (Sigma Aldrich) were immobilized on all channels of a CM5 sensor chip. Approximately 200 RU of Fc-tagged IL-1RAcP were directly captured from cell-culture supernatants on flow cell 2, 3, or 4. Binding experiments were carried out in 10 mM HEPES, pH 7.4, 150 mM NaCl, 0.05% (v/v) Tween20 at 25C by single cycle kinetic analysis using five concentrations of CAN03 or CAN04 Fab, from 20nM to 1.25nM concentrations. Between runs, the sensor surface was regenerated with one 45s injection of 10 mM HCl. Sensorgrams were double referenced against the control flow cell and buffer injections. Data were fit to a 1:1 binding model using the Biacore T100 Evaluation Software. In reference to Günther *et al*., numbering of alanine scan residues is offset by 20 to reflect human IL1β/IL-1RI/IL-1RAcP crystal structure (pdb:4DEP) [22, 30].

### Bi-Epitopic Inhibition Assays

To measure the potency of the antibodies, HEK-Blue™(IL-1β/IL-33) cells were seeded in a 96 well plate at a concentration of 0.33 × 10^6^ cells/ml. Nine 1:3 serial dilutions of antibodies from a stock of 1uM antagonist were added to the cells and incubated for 45 min for a final concentration of 200, 66.7, 20, 6.7, 2, 0.67, 0.2, 0.067, 0.02, 0.0067 nM. Plates were stimulated with 10ul of cytokine (1ng/mL IL-1β or 3ng/mL IL-33 stocks at 20X) and incubated for 18hr at 37C. Cells were then harvested and analyzed using the QuantiBlue assay. As a negative control, activity without cytokine stimulation was measured; as a positive control, cells were stimulated without antibody addition.

## Results

### CAN03 and CAN04 both exhibit high affinity binding to their common IL-1RAcP antigen

Antibody affinity has long been known to play a fundamental role in governing the biological consequence of antibody/antigen interactions. To determine the binding parameters of the antibodies CAN03 and CAN04 to IL-1RAcP, we conducted surface plasmon resonance (SPR) analysis. Both CAN03 and CAN04 display a high affinity for their target IL-1RAcP, exhibiting affinities (K_D_) of 900pM and 780pM, respectively, and slow off-rates (K_d_) near 5 × 10^−4^ 1/s (Figure 1A and 1B). The on-rates (K_a_) of both CAN03 and CAN04 are similar as well, at 7.90 × 10^5^ (1/Ms) and 6.10 × 10^5^ (1/Ms), respectively. To determine if native glycosylation of IL-1RAcP, on which there are seven putative glycosylation sites, influenced binding, we generated a high-mannose variant of IL-1RAcP and subsequently cleaved off the glycans using an endoglycosidase [22]. The previous SPR experiments were then repeated using de-glycosylated IL-1RAcP as the antigen. For both CAN03 and CAN04 antibodies, there was no appreciable difference in affinities for complexes formed with the natively glycosylated or deglycosylated antigen IL-1RAcP (Figure 1C and 1D). This lack of change demonstrates that epitope masking by glycosylation does not occur as glycosylation state does not affect binding, as would be seen by an increase in affinity. In addition, the glycans of the native protein do not constitute part of the binding epitope for either antibody, as would be seen by a decrease in affinity by their removal.

**Figure 1:**
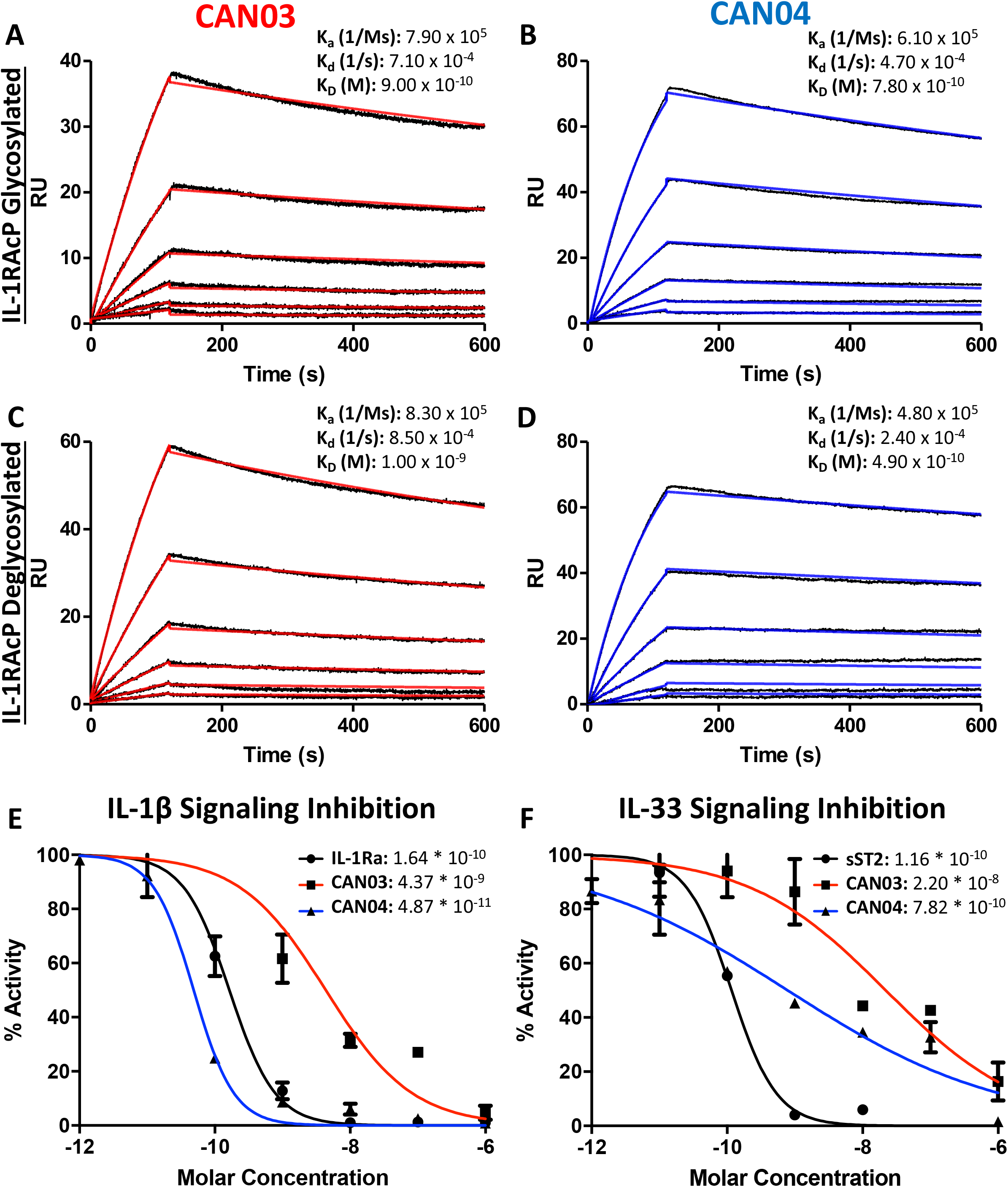
Surface Plasmon Resonance of the antibodies CAN03 & CAN04 and Cell Signaling Assays. **(A)** Sensorgram of CAN03 with natively glycosylated IL-1RAcP as the antigen and the K_a_, K_d_, and K_D_ of interaction labelled (Curve fit: red). **(B)** Sensorgram of CAN04 with natively glycosylated IL-1RAcP as the antigen and the K_a_, K_d_, and K_D_ of interaction labelled (Curve fit: blue). **(C)** Sensorgram of CAN03 with deglycosylated IL-1RAcP as the antigen and the K_a_, K_d_, and K_D_ of interaction labelled (Curve fit: red). **(D)** Sensorgram of CAN04 with deglycosylated IL-1RAcP as the antigen and the K_a_, K_d_, and K_D_ of interaction labelled (Curve fit: blue). **(E)** Cell signaling assays of IL-1 with the natural antagonist IL-1Ra, anti-IL-1RAcP antibody CAN03, and anti-IL-1RAcP antibody CAN04. IC_50_ values are labelled. **(F)** Cell signaling assay of IL-33 with the natural antagonist sST2, anti-IL-1RAcP antibody CAN03, and anti-IL-1RAcP antibody CAN04. IC_50_ values are labelled.

### CAN03 and CAN04 inhibit IL-1β with different potencies but both preferentially inhibit IL-1β over IL-33

Next, we investigated the effects that CAN03 and CAN04 have on IL-1β and IL-33 signaling. As IL-1β and IL-33 signaling complexes interact with IL-1RAcP differently [22], we surmised that antibodies that target different epitopes on IL-1RAcP would affect IL-1 and IL-33 signaling differently. As a standard of cytokine signaling inhibition, we used the natural inhibitors to IL-1 and IL-33 signaling, IL-1 receptor antagonist (IL-1Ra), known pharmacologically as Anakinra, and soluble ST2 (sST2), respectively. Unlike the antagonist cytokine IL-1Ra, sST2 is the soluble isoform of ST2 and acts as a decoy receptor to antagonize IL-33 signaling [31]. CAN04 was the most potent inhibitor of IL-1β signaling, exhibiting an IC_50_ of 49 pM in comparison to the IC_50_ of 164 pM for the antagonist cytokine IL-1Ra (Figure 1E). Conversely, CAN03 was the least potent inhibitor of IL-1β. CAN03 was a 90-fold weaker inhibitor of IL-1β signaling, with an IC_50_ of 4.4 nM, than was CAN04 (Figure 1E). CAN04 was also a stronger inhibitor of IL-33 than CAN03, exhibiting an IC_50_ of 782pm compared to an IC_50_ of 22nM for CAN03 (Figure 1F). Neither CAN03 nor CAN04 antibodies inhibited IL-33 as well as the natural antagonist soluble ST2 (sST2), however, as sST2 displayed an IC_50_ of 116pM. (Figure 1F). The antibody CAN03 was the least potent inhibitor in both cases, displaying a 25-fold worse IC_50_ against IL-1β signaling than IL-1Ra and a 190-fold worse IC_50_ against IL-33 signaling than sST2 (Figure 1E and 1F).

### CAN03 and CAN04 use different combinations of CDRs to engage distinct epitopes on IL-1RAcP

To determine the complementary-determining regions (CDRs) of the antibodies that comprise the binding interface with IL-1RAcP, we conducted hydrogen deuterium exchange coupled with mass spectrometry (HDX-MS). First, we measured deuterium uptake of CAN03 and CAN04 in their apo forms. When compared with deuterium exchange profiles of their antigen bound forms, areas of protection, imparted by conformational restriction and loss of solvent accessibility to antigen bound areas, are seen as a decrease in deuterium exchange (Figure 2A and 2B). For the CAN03 Fab, both CDR1 and CDR2 of the heavy chain are protected upon binding to IL-1RAcP, as seen on peptide fragments 27-32 and 51-57, respectively (Figure 2C). On the light chain, CDR2 and CDR3 are protected, as seen on peptide fragments 47-53 and 93-106 (Figure 2C). For the antibody CAN04, the heavy chain utilizes all three of its CDRs in IL-1RAcP binding, as seen on peptide fragments 24-29, 47-69, and 98-103 (Figure 2D). On the light chain, only CDR2 displays binding, as seen on peptide fragment 47-54 (Figure 2D).

**Figure 2:**
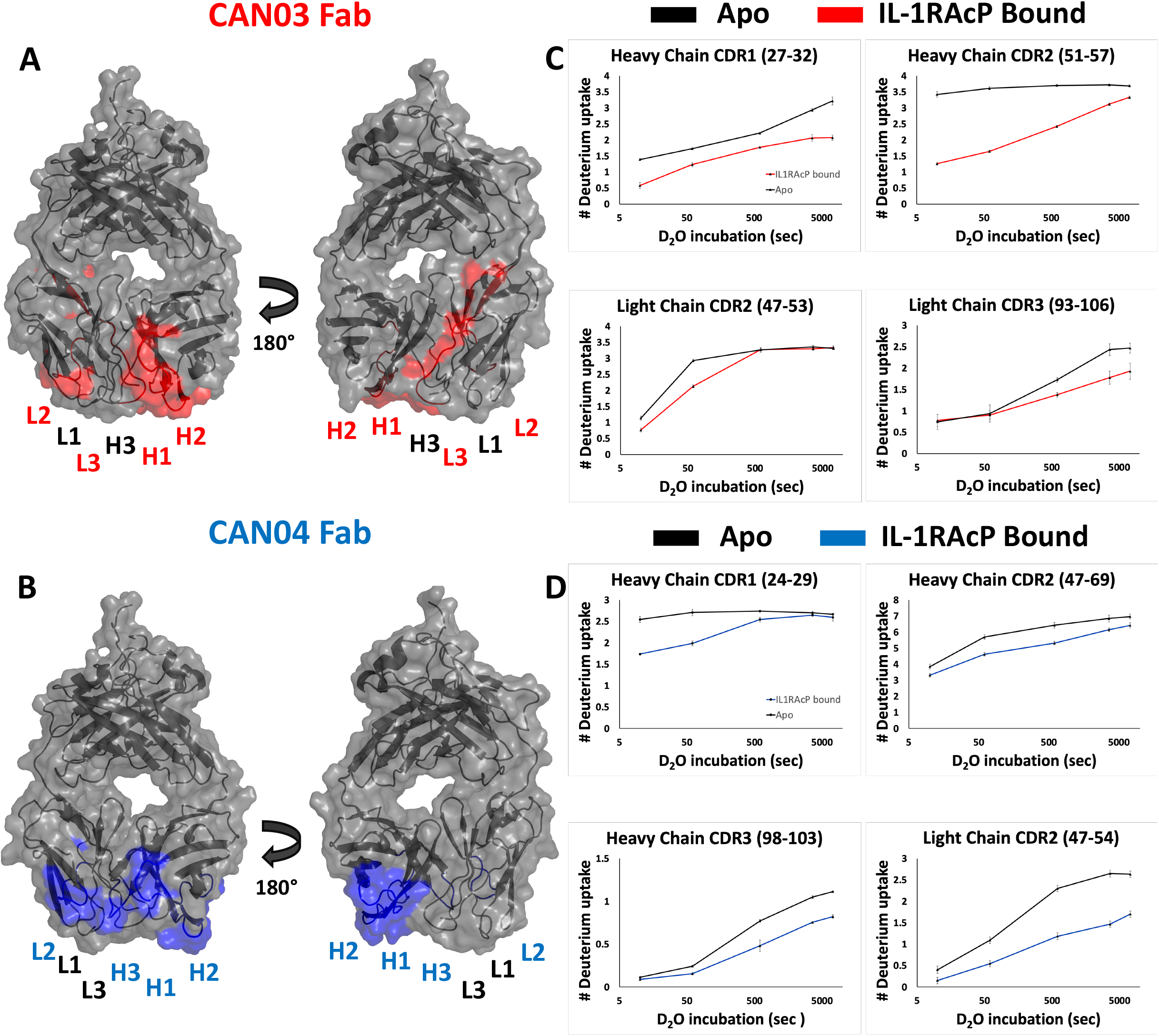
Deuterium exchange protection on the antibodies CAN03 and CAN04 by IL-1RAcP. **(A)** Model of Fab CAN03 with binding by HDX-MS colored red. **(B)** Model of Fab CAN04 with binding by HDX-MS colored blue. **(C)** Peptide stretches identified by HDX-MS for deuterium exchange differences between IL-1RAcP bound CAN03 and Apo form CAN03. **(D)** Peptide stretches identified by HDX-MS for deuterium exchange differences between IL-1RAcP bound CAN04 and Apo form CAN04.

To ascertain the molecular mechanism of selective inhibition of IL-1 over IL-33 signaling, we sought to determine the epitopes on IL-1RAcP that CAN03 and CAN04 recognize. Again using HDX-MS analysis, we found CAN03 and CAN04 antibodies bind distinct domains of IL-1RAcP (Figure 3A and 3B). CAN03 binds to domain 3 (D3) of IL-1RAcP, which is proximal to the plasma membrane (Figure 3A). This binding occurs along four sequential peptide fragments that span residues 254-296 (Supplemental Figure 1A). In the first peptide fragment (254-259), there is a clear reduction of deuterium exchange by CAN03 bound to IL-1RAcP compared to the exchange that occurs by IL-1RAcP in its unbound, apo state (Supplemental Figure 1A). This exchange profile continues in the peptide fragments 259-275 and 276-288. In the last fragment, 290-296, the exchange profile is not as pronounced. As there is not a large difference at time-point 10 sec, the exchange seen may be due to restricted movement within this stretch of amino acids by direct binding in the proceeding regions. Notably, only IL-1RAcP peptides residing entirely within the D3 domain, and not in the D1 or D2 domains, are involved in binding CAN03. Conversely, CAN04 binds domain 2 (D2) of IL-1RAcP (Figure 3B). The protection imparted by CAN04 binding IL-1RAcP is seen on the three peptides comprising residues 105-114, 145-158, and, 169-176 (Supplemental Figure 1B). While these peptides are not sequential, they reside adjacent to one another in the folded protein on the top of D2 (Figure 3B).

**Figure 3:**
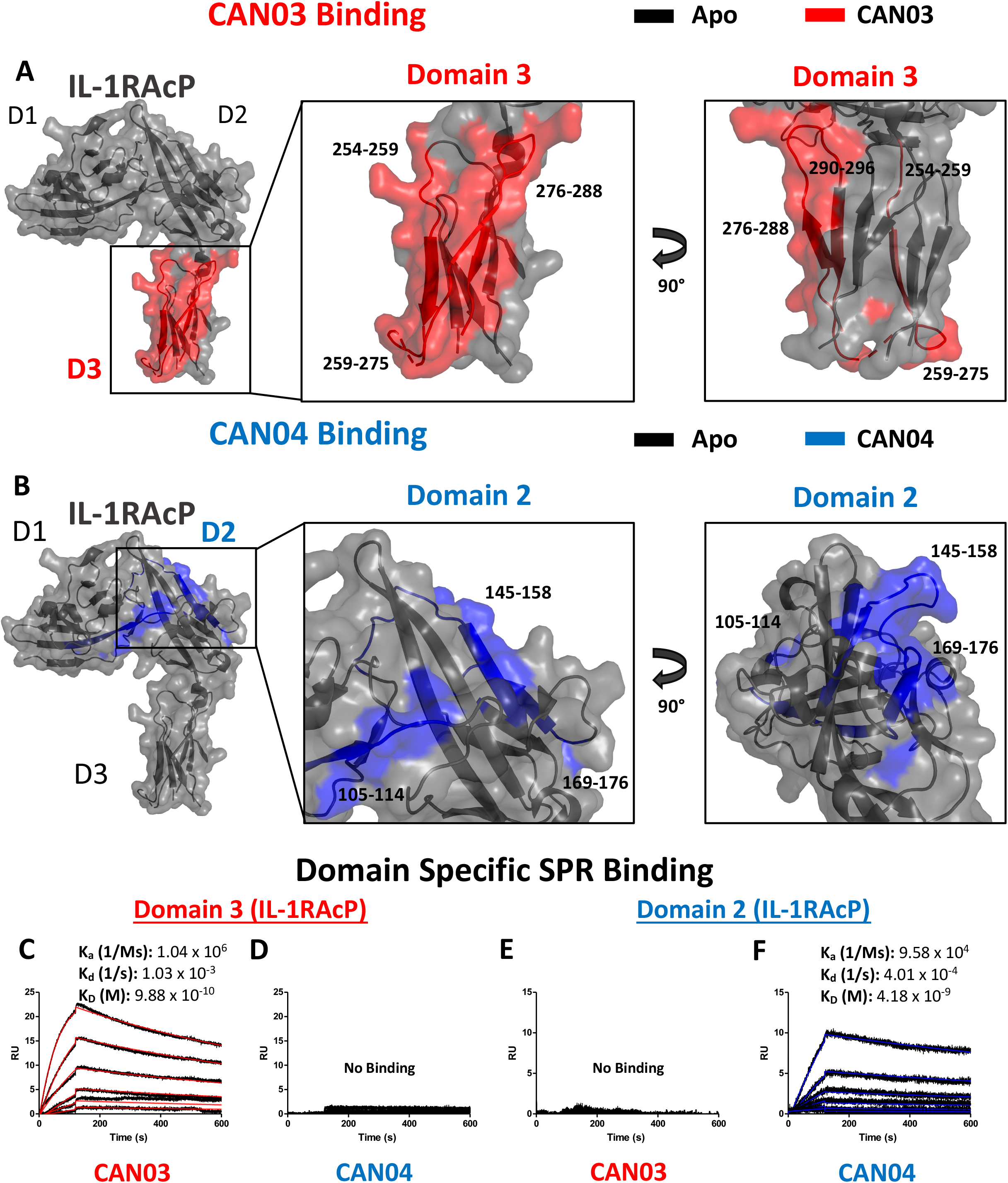
Deuterium exchange protection on IL-1RAcP by the antibodies CAN03 and CAN04. **(A)** Crystal structure of IL-1RAcP (pdb:4DEP) with CAN03 binding by HDX-MS colored red. Domain 3 is englarged to show locations of the four peptide regions 254-259, 259-275, 276-288, and 290-296 from both a front and side view. **(B).** Crystal structure of IL-1RAcP (pdb:4DEP) with CAN04 binding by HDX-MS colored blue. Domain 2 is englarged to show locations of the four peptide regions 105-114, 145-158, and 169-176 from both a front and side view. **(C)** Sensorgram of CAN03 with IL-1RAcP Domain 3 as the antigen and the K_a_, K_d_, and K_D_ of interaction labelled (Curve fit: red). **(D)** Sensorgram of CAN04 with IL-1RAcP Domain 3 as the antigen (no binding). **(E)** Sensorgram of CAN03 with IL-1RAcP Domain 1&2 as the antigen and kinetics of interaction labelled (no binding). **(F)** Sensorgram of CAN04 with IL-1RAcP Domain 1&2 as the antigen and the K_a_, K_d_, and K_D_ of interaction labelled (Curve fit: blue).

To verify that CAN03 and CAN04 have distinct and non-overlapping epitopes on IL-1RAcP, we produced two fragments of IL-1RAcP, containing either a combination of the D1 and D2 domains (D1/2) or D3 domain alone, and measured the binding parameters of the CAN03 and CAN04 antibodies to these IL-1RAcP fragments by SPR. CAN03 binds specifically to the D3 domain of IL-1RAcP with approximately the same affinity as it did for full-length IL-1RAcP (Figure 3C). CAN04, however, shows no appreciable binding for D3 by SPR (Figure 3D). Conversely, CAN04 binds to the D1/2 domains of IL-1RAcP, albeit with a weaker affinity than the full length, while CAN03 displays no binding to this region, providing further evidence that these antibodies bind distinct regions of their common antigen, IL-1RAcP (Figure 3E and 3F).

### Molecular modeling of antibody-IL-1RAcP complexes and Chimeric IL-1RAcP Binding

To visualize possible binding modes of the antibodies CAN03 and CAN04, we conducted molecular modeling of the antibody-IL-1RAcP complexes. Using our Fab constructions in combination with our HDX-MS and SPR data as constraints, we generated multiple poses for CAN03 and CAN04 bound to IL-1RAcP. Our models indicate that CAN03 binds domain 3, near yet not overlapping with the binding interface of IL-1β, while CAN04 binds domain 2, directly coincident with portions of the IL-1β and IL-33 signaling complexes. (Figure 4A and 4B). We further validated these poses by comparing the modeled change in deuterium uptake versus our experimental deuterium uptake by IL-1RAcP. Both CAN03 and CAN04 poses aligned well with our empirical data, displaying an root-mean-square deviation (RMSD) of deuterium exchange of 0.981 Åfor CAN03-IL-1RAcP and 0.684 Åfor CAN04-IL-1RAcP (Figure 4C and 4D).

**Figure 4:**
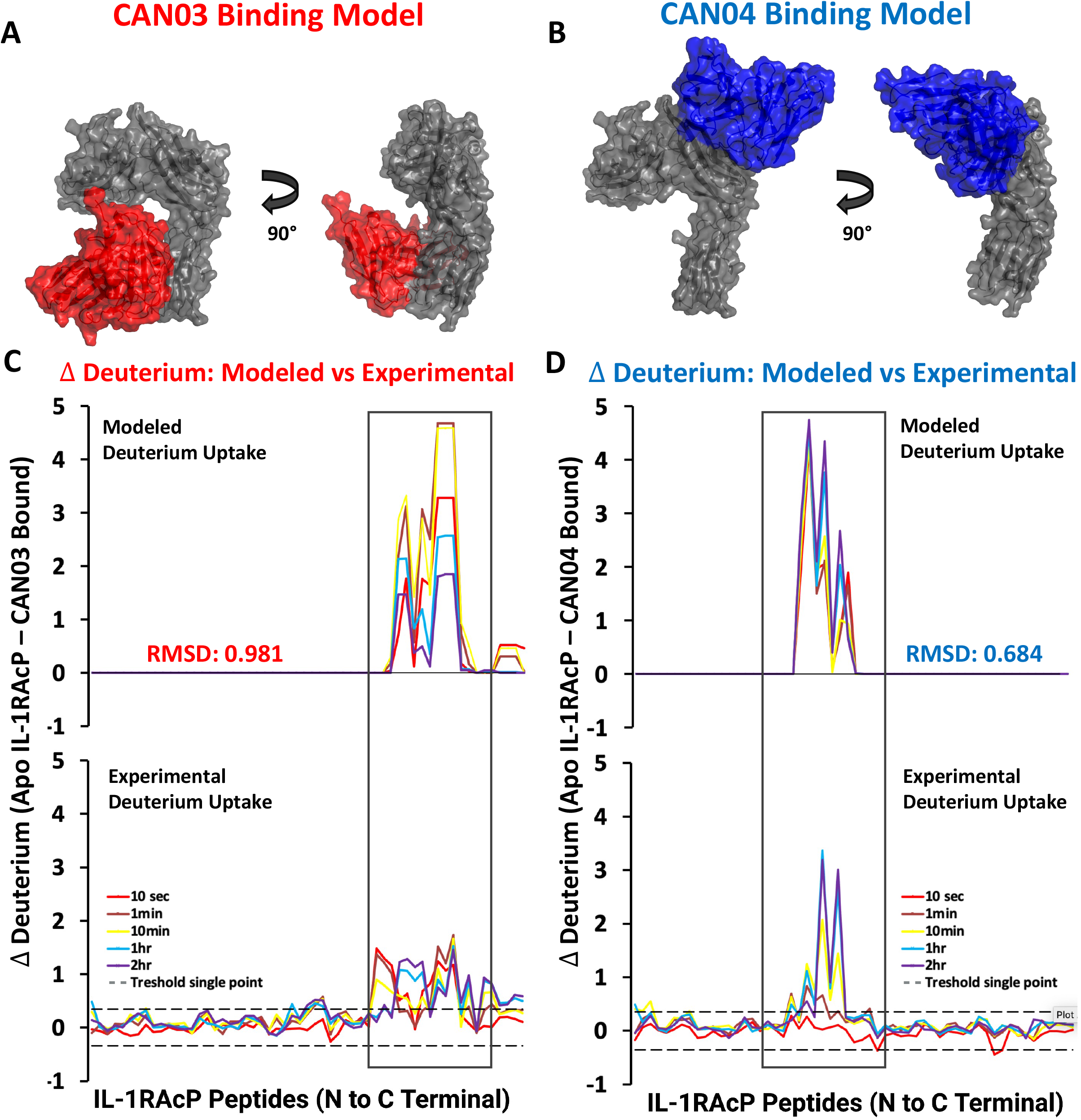
Models CAN03/IL-1RAcP and CAN04/IL-1RAcP binding and deuterium uptake comparison. **(A)** Model of CAN03 Fv (Red) binding IL-1RAcP. **(B)** Model of CAN04 Fv (blue) binding IL-1RAcP. **(C)** Difference plot of modelled deuterium uptake versus experimental deuterium uptake for CAN03 (Red). **(D)** Difference plot of modelled deuterium uptake versus experimental deuterium uptake for CAN04 (Blue).

To experimentally test our structural models and to further clarify the binding region of CAN04, we generated chimeras of IL-1RAcP with portions of the human IL-1RAcP (hIL-1RAcP) in a murine IL-1RAcP (mIL-1RAcP) background. In all, we generated three chimeric proteins containing two regions of hIL-1RAcP within D2 (Figure 5A and 5B). As expected, CAN04 binds hIL-1RAcP; conversely, CAN04 does not bind mIL-1RAcP (Figure 5C). When we added human region 1 (H1; incusive of residues Pro121 to Arg137) and human region 2 (H2; inclusive of residues Thr154 to Ile171) to mIL-1RAcP, we observed restored binding by CAN04. When only H1 was included without H2, we observed ablated CAN04 binding. If H2 was added to mIL-1RAcP, even in the absence of H1, binding of CAN04 was re-established, displaying the necessity of H2 for CAN04 binding, an area implicated in our HDX-MS data, modelling data, and which contains the c2d2 loop (Figure 5C).

**Figure 5:**
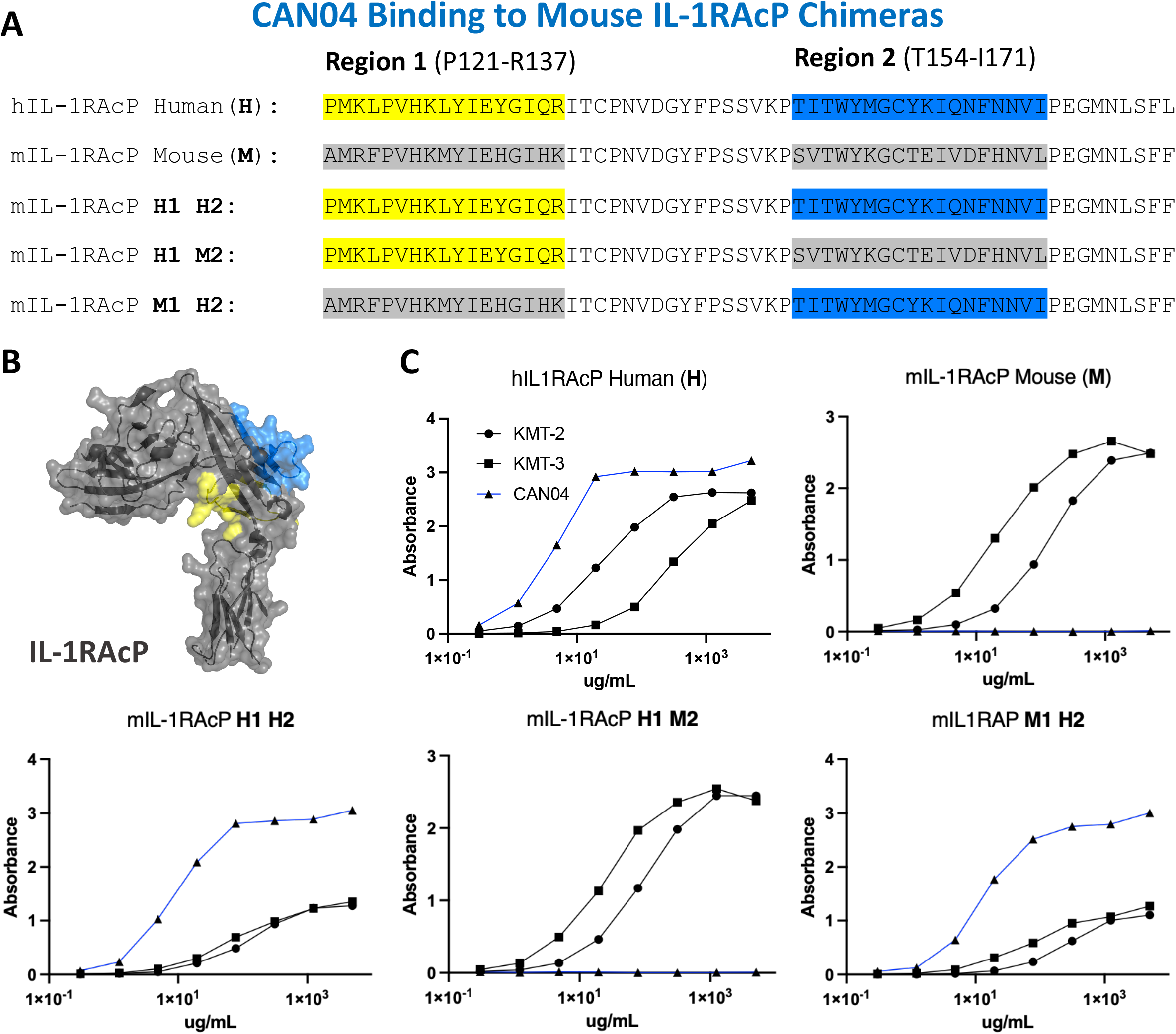
CAN04 binding to mouse IL-1RAcP chimeras. **(A)** Alignment of human IL-1RAcP, mouse IL-1RAcP, and chimeric mIL-1RAcP with human regions 1 and/or 2 colored. **(B)** Crystal structure of human IL-1RAcP (pdb: 4DEP) with regions 1 and 2 colored **(C)**. Binding profile of CAN04 (mAb) to each version of IL-1RAcP) with KMT-2 (polyclonal hIL-1RAcP antibody mixture) and KMT-3 (polyclonal mIL-1RAcP antibody mixture) as controls.

### Structural analysis and alanine scan elucidates mechanism of selective inhibition of cytokine signaling

To further probe the interfaces of CAN03 and CAN04 on IL-1RAcP, we conducted an analysis of the HDX-MS data in conjunction with the crystal structures of IL-1β (pdb:4DEP) and IL-33 (pdb:5VI4) signaling complexes using PISA and known interface residue energy contributions for each signaling complex (Figure 6A and 6B) [22, 30, 32]. After mapping the interface between IL-1RAcP and the IL-1 and IL-33 cytokine/receptor pairs, we mutated residues that overlapped or were near CAN03 and CAN04 binding areas as seen by HDX-MS (Figure 6C and 6D). By measuring the binding energy change, ΔΔG, of each mutant compared to wild-type IL-1RAcP, we determined the energetic contribution of each IL-1RAcP interface residue for CAN03 and CAN04 binding (Figure 6E).

**Figure 6:**
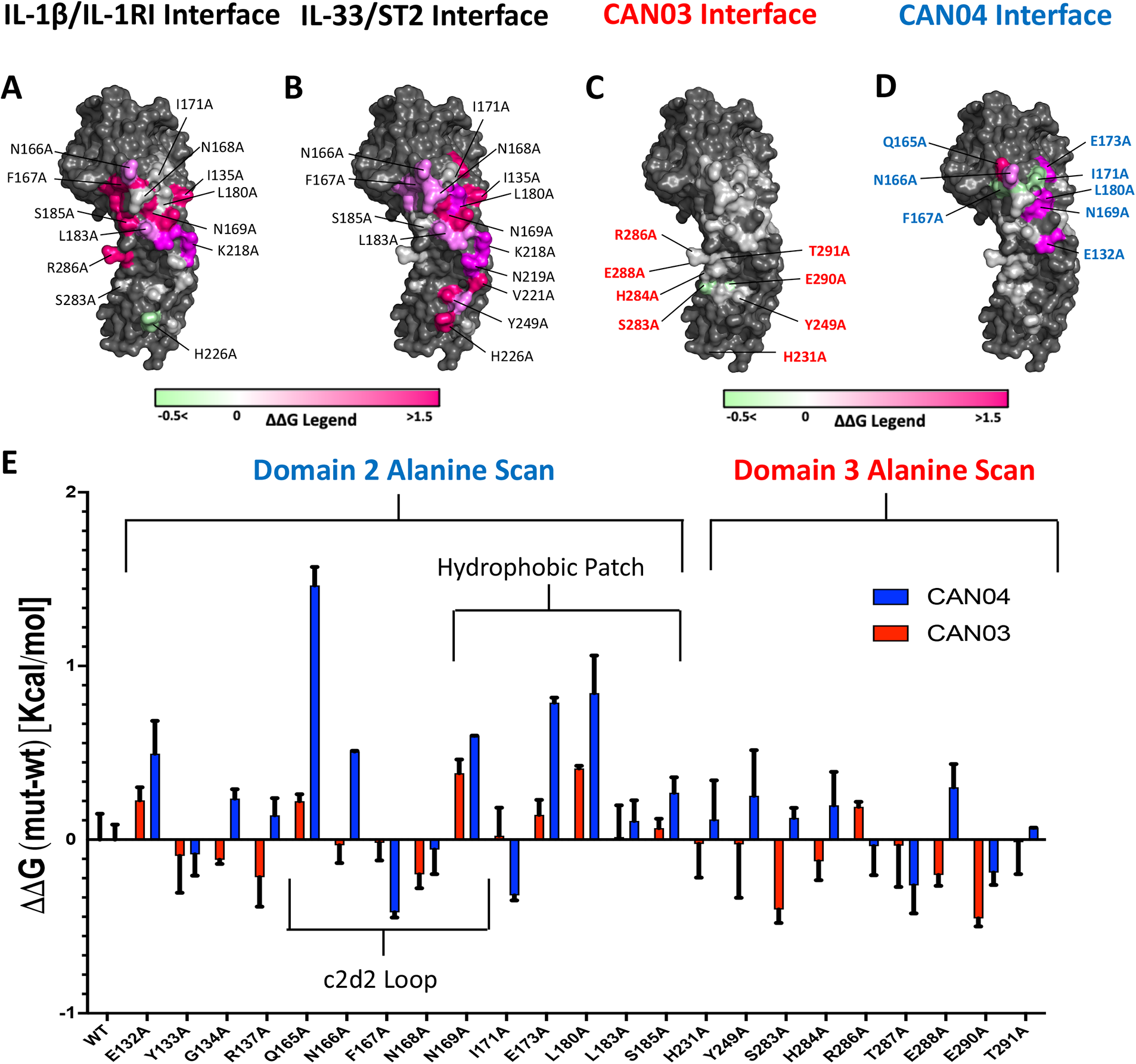
Extensive Alanine Scan of IL-1RAcP by Surface Plasmon Resonance. **(A)** IL-1RAcP with IL-1*β*/IL-1RI interface colored according to known ΔΔG of contributing residues **(B)** IL-1RAcP with IL-33/ST2 interface colored according to known ΔΔG of contributing residues **(C)** IL-1RAcP with CAN03 interface colored according to experimental ΔΔG of contributing residues **(D)** IL-1RAcP with CAN03 interface colored according to experimental ΔΔG of contributing residues **(E)** Graph of ΔΔG (mut-wt) of CAN03 and CAN04 Fabs to IL-alanine library of IL-1RAcP.

In our PISA analysis, clear differences arise in D3 utilization by IL-1β and IL-33 signaling complexes [32]. Residues in the interface of D3 contribute only 231 Å^2^ of buried surface area (BSA) in IL-1β complex formation, as opposed to 934 Å^2^ BSA of D2 and 137 Å^2^ BSA of the linker. This translates to fewer residues in D3 being involved in IL-1β/IL-1RI recruitment than D2 and the linker. Although we included additional residues important for IL-1RAcP signaling, namely H231A and Y249A, the vast majority of the interface in D3 of IL-1/βIL-1RI is contained on the peptide stretch from S283 to T291. While we observed a moderate negative energy change in residue S283 and E290, near -0.4 kcal/mol, there was little change in ΔΔG over the entirety of this span of residues (Figure 6E).

For IL-33 signaling, the D3 domain has nearly as large an interface area as D2, comprising 704 Å^2^ BSA to IL-33/ST2 in comparison to 923 Å^2^ for D2 and 139 Å^2^ for the linker. The majority of residues involved in IL-33/ST2 binding, however, are far outside of the HDX-MS binding stretches, namely from V221-V232, due to a 60 degree turn of D3 in the IL-33 crystal structure compared to IL-1β [22]. Y249A is a part of the interface, although we saw little change in our alanine scan of this residue (Figure 6E).

For the antibody CAN04, the mutation Q165A decreased the affinity of CAN04 the greatest in our alanine scan, by approximately 1.5 kcal/mol, and likely is a critical part of the binding interface with CAN04 (Figure 6F). Directly beside this residue, the mutation N166A decreased binding by 0.5 kcal/mol while F167A increased binding by -0.4 kcal/mol. Adjacent to this residue, N169A also affected ΔΔG of CAN04 binding by roughly 0.6 kcal/mol. In addition to this stretch of residues, the residues E173 and L180 contributed nearly 1 kcal/mol of energy to the CAN04 interaction (Figure 6F). Proximal to these residues, I171A increased binding by 0.3 kcal/mol.

### Potency of signaling inhibition by CAN03 and CAN04 is increased in a bi-epitopic antibody format

Since CAN03 and CAN04 have distinct binding interfaces on different domains of IL-1RAcP, we created a bi-epitopic library of the antibodies to determine if the combination of CAN03 and CAN04 would have a synergistic effect against IL-1β and IL-33 signaling (Figure 7A). In addition to the canonical IgG forms described above, we tested three different antibodies: a CAN03 tetravalent antibody with CAN04 (tetra-CAN03-CAN04; in which the CAN04 Fab is appended to the CAN03 antibody), a CAN04 tetravalent antibody with CAN03 (tetra-CAN04-CAN03; in which the CAN03 Fab is appended to the CAN04 antibody), and a bispecific antibody that contains one of each CAN03 and CAN04 Fab (Figure 7A). Neither Tetra-CAN03-CAN04 nor Tetra-CAN04-CAN03 were significantly more potent as inhibitors against IL-1β or IL-33 signaling than either CAN03 or CAN04 alone (Figure 7B). The bispecific antibody, however, inhibited both IL-1β and IL-33 better than both CAN03 and CAN04 alone (Figure 7B and 7C). As the IL-1RAcP epitope for CAN04 was already optimally positioned for potent IL-1β inhibition, the improvement seen by the inclusion of CAN03 in the bi-specific antibody was modest. The bi-specific antibody exhibited the greatest effect against IL-33 signaling, acting synergistically to block IL-33 better than both CAN03 and CAN04 alone, as each antibody could block both the D3 and D2 simultaneously on IL-1RAcP (Figure 7C). As we used different IL-1β and IL-33 signaling cells for these studies, different concentrations of both IL-1β and IL-33 were required in our experimental design; as such, only the relative inhibitory concentrations of our antagonists (Figure 7B-C) are directly comparable to the signaling assays described previously (Figure 1E-F).

**Figure 7:**
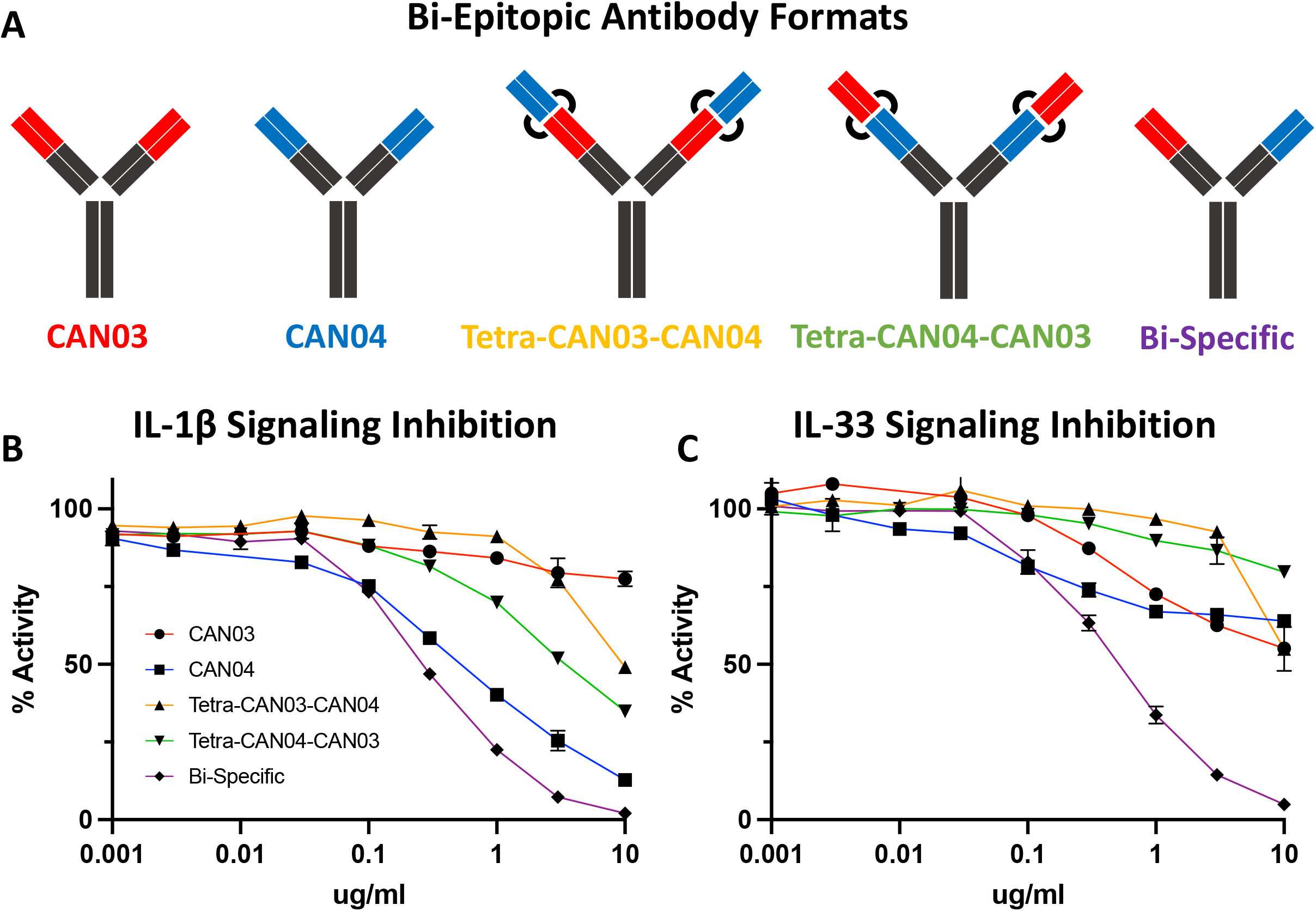
Bi-Epitopic Antibodies and Signaling Inhibition Studies. **(A)** Library of antagonists: CAN03 mAb, CAN04 mAb, tetravalent CAN03 with CAN04 attached to Fab (Tetra-CAN03-CAN04), tetravalent CAN04 with CAN03 attached to Fab (Tetra-CAN04-CAN03), and bi-specific CAN03/CAN04 (Bi-Specific) antibody. **(B)** Signaling inhibition assay of IL-1 including all five antagonists with identities colored and labelled. **(C)** Signaling inhibition assay of IL-33 including all five antagonists with identities colored.

## Discussion

In many respects, the IL-1β and IL-33 signaling complexes are similar. First, the cytokine binds its cognate receptor at high (sub-nM to low nM) affinity; second, the cytokine/cognate receptor complex recruits the shared co-receptor IL-1RAcP at more moderate (high nM) affinity. IL-1RAcP interacts with both IL-1β/IL-RI and IL-33/ST2 binary complexes similarly, using its D2 and D3 domains. There are four main surface regions on IL-1RAcP that contribute to these interactions: the c2d2 loop (Q165-N169), the hydrophobic patch (E132-I135, I171-S185), the linker between D2 and D3 (G215-N219), and the D3 domain (V221-T291) [22, 30]. The respective crystal structures of the two ternary complexes are highly similar and differ only by a root-mean-square deviation of 3.2 Å over all Cα carbons [33]. The differences, however, reside in how the respective cytokine/cognate receptor binary complexes utilize the four surface regions of IL-1RAcP in order to form functional signaling complexes.

In IL-1 signaling, the vast majority of the IL-1β/IL-1RI interface is contained within domain 2 of IL-1RAcP. The c2d2 loop, inclusive of residues Q165 to N169, is crucial to this interaction. Indeed, Q165 of IL-1RAcP hydrogen bonds with Q141 of IL-1β while not making any crystal contacts in the IL-33 signaling complex. F167 of IL-1RAcP contributes eight-fold more energetically to the complex formation of IL-1 than it does to IL-33, translating to 4 kcal/mol in IL-1β/IL-1RI binding energy (Figure 6C and D) [22]. Adjacent to this residue, N169 contributes even more binding energy, 4.5 kcal/mol, to IL-1β/IL-1RI binding, two-fold more than in the IL-33/ST2 complex [22]. Within the entirety of this stretch of residues, a network of Van der Waals interactions, hydrogen bonds, and electrostatic contacts exist.

The hydrophobic patch is an important component to IL-1β complex formation as well. Energetically, E132 is two-fold more important to IL-1β signaling than IL-33, translating to 2 kcal/mol. S185 contributes nearly six-fold more to IL-1β complex formation than IL-33, adding 3 kcal/mol binding energy to the interaction [22]. G134 hydrogen bonds with IL-1RI; I171 and L180 each are part of a hydrophobic interaction with V160 of IL-1RI. D3, however, is less utilized than D2, and the entirety of tested residue interactions only contribute 3 kcal/mol of binding energy to IL-1 complex formation. These interactions are mainly localized between a stretch of amino acids from S283 and T291, although H231 and Y249 make minor energetic contributions [22].

CAN03 is a weaker inhibitor of IL-1β signaling than IL-1Ra and CAN04 for two reasons. First, D3 is not as energetically important for IL-1 signaling as D2 and is therefore a less direct target for inhibiting IL-1β signaling (Figure 6A). Second, as shown by our alanine scan, CAN03 does not directly interact with residues involved in IL-1β signaling and likely binds residues prior in sequence to S283-T291 (Figure 6C). Altogether, the limited IL-1β signaling inhibition by CAN03 is most likely due to steric effects resulting from the bulkiness of the IgG molecule rather than direct binding of the interface important for IL-1β complex formation.

CAN04 is a potent inhibitor of IL-1β signaling for the very reason CAN03 is not: CAN04 binds directly to residues that are integral to IL-1β signaling. This is seen in our chimeric binding studies, wherein region 2 (T154-I171), an area containing the c2d2 loop, was essential for CAN04 binding and its absence resulted in loss of binding. Addtionally, in our alanine scan, the largest energy change seen was for Q165A, a residue contributing 1.5 kcal/mol energy to CAN04 binding. N166A, F167A, and N169A all affected CAN04 binding as well (Figure 6F). Collectively, these residues comprise the c2d2 loop and are clearly part of the CAN04 binding interface. Near this area, we measured large energetic changes for the E173A and L180A mutants. Altogether, CAN04 binds the c2d2 loop and, to a lesser extent, portions of the hydrophobic patch, to selectively inhibit IL-1β signaling.

While IL-1 signaling is highly dependent on the c2d2 loop and hydrophobic patch, recruitment of IL-1RAcP by IL-33/ST2 is the result of a more distributed interface. This is due to IL-33 holding ST2 in a specific conformation for IL-1RAcP recruitment rather than IL-33 engaging with IL-1RAcP to a similar extent as does IL-1β. As a consequence, the c2d2 loop is utilized less [22]. Within the hydrophobic patch, however, E173 and L180 contribute the most energy, roughly 1.5 kcal/mol, to IL-33/ST2 binding. Overall, D2 is not as energetically important in IL-33 signaling as in IL-1β signaling. D3, in contrast to its role in IL-1β signaling, is critically important for IL-33 signaling. Within the IL-33 crystal structure, D3 is rotated 64 degrees in comparison to its position in the IL-1β signaling complex. As Ig domains are ellipsoidal, this results in a larger surface area being presented to IL-33/ST2 in D3 of IL-1RAcP and along a different stretch of residues, namely V221–V232 and Y249 [22].

While the IL-1β interface is near the epitope of CAN03, the IL-33 interface is not, and most likely accounts for the 190-fold worse IL-33 signaling inhibition seen in our assays as compared to sST2 (Figure 1F). This is highlighted best in our HDX-MS data, where CAN03 binding is clearly to the backside of the D3 in relation to the V221-V232 interface (Figure 3A). In further iterations of IL-1RAcP antibody design, it may be possible to selectively inhibit IL-33 over IL-1 signaling by targeting these residues of IL-1RAcP as there is little overlap between the IL-1 and IL-33 interface in the D3 domain (Figure 6A and 6B).

The shortcomings of CAN04 as an IL-33 inhibitor are two fold. First, the c2d2 loop does not contribute to IL-33 complex formation to the same degree as it does for IL-1β. Additionally, as the IL-33/ST2 interaction is more distributed on the IL-1RAcP surface and relies heavily on D3, the CAN04 binding epitope is poorly positioned to be nearly as potent an IL-33 inhibitor as it is an IL-1 inhibitor. In our bi-epitopic antibody design, it is not entirely surprising the bi-specific CAN03/CAN04 antibody performed better than either CAN03 and CAN04 alone against IL-33 signaling (Figure 7C). As the interface between IL-1RAcP and IL-33/ST2 is broad, targeting both D2 and D3 simulataneoulsy with the bi-specific antibody appears to be a powerful strategy for IL-33 signaling inhibition.

Collectively, our studies highlight the feasibility of using antibodies to target shared secondary receptors for selective cytokine signaling inhibition of IL-1 family cytokines. By logical extension, the antibodies could be replaced by other macromolecules that bind similarly specific interfaces and the IL-1 family cytokines substituted by other cytokine families. Indeed, shared receptors abound in cytokine signaling. Within the class I cytokine receptor family alone, three shared receptors, the common gamma chain (γ_c_), gp130, and the common beta chain (β_c_), are collectively involved in nearly 20 different cytokine complexes [34]. Within the class II cytokine receptor family, four shared receptors are involved in 9 different cytokine complexes [35].

Through targeting a shared receptor and leveraging the differential utilization of shared interfaces, it is possible to selectively inhibit one cytokine over another, albeit with different efficacy. In addition, we identified an antibody that inhibits IL-1 signaling better than the natural antagonist cytokine, IL-1Ra (Anakinra). As Anakinra is the gold-standard of IL-1 therapeutics, CAN04 may prove clinically useful for a wide range of IL-1-mediated inflammatory and autoimmune diseases.

## Acknowledgements

This work was supported by NIH grants AI132766, AI132766-02S1 (to EJS), and Cantargia AB.

## Notes

### Competing Interest Statement

Competing Interests Statement: D.L, G.F., K.S., and G.B. are employees of Cantargia AB (Medicon Village, Lund, Sweden). Cantargia AB is the owner of the intellectual property rights for CAN03 and CAN04 for use in the treatment and diagnosis of neoplastic disorders. Cantargia AB partially funded this research. The remaining authors declare no competing financial interests.

