## Supplemental Text for "Molecular basis of selective cytokine signaling inhibition by antibodies targeting a shared receptor"

**Supplemental Figure 1: Deuterium uptake of IL-1RAcP in apo form and antibody bound form. (A)** Deuterium uptake graphs of CAN03 bound (red), CAN04 bound (blue), and Apo IL-1RAcP. This stretch of residues was used in figure 4A for our HDX-MS analysis of CAN03 binding to IL-1RAcP. **(B)** Deuterium uptake graphs of CAN03 bound (red), CAN04 bound (blue), and Apo IL-1RAcP. This stretch of residues was used in figure 4B for our HDX-MS analysis of CAN04 binding to IL-1RAcP.

**Supplemental Figure 2: Sensograms from corresponding SPR experiments for Extensive Alanine Scan (A)** Sensograms of IL-1RAcP alanine scan (Fc-fused IL-1RAcP with alanine mutations) and CAN03 as the analyte. **(B)** Sensograms of IL-1RAcP alanine scan (Fc-fused IL-1RAcP with alanine mutations) and CAN04 as the analyte.
