## Supplementary figures and images for "Molecular basis of selective cytokine signaling inhibition by antibodies targeting a shared receptor"

### Supplemental Figures

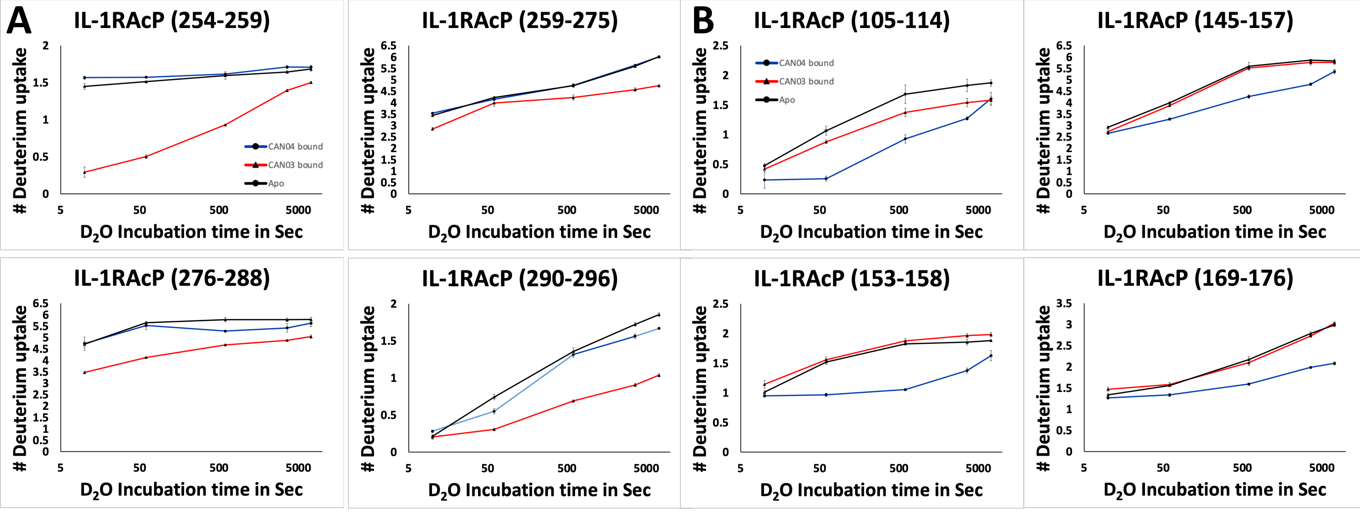

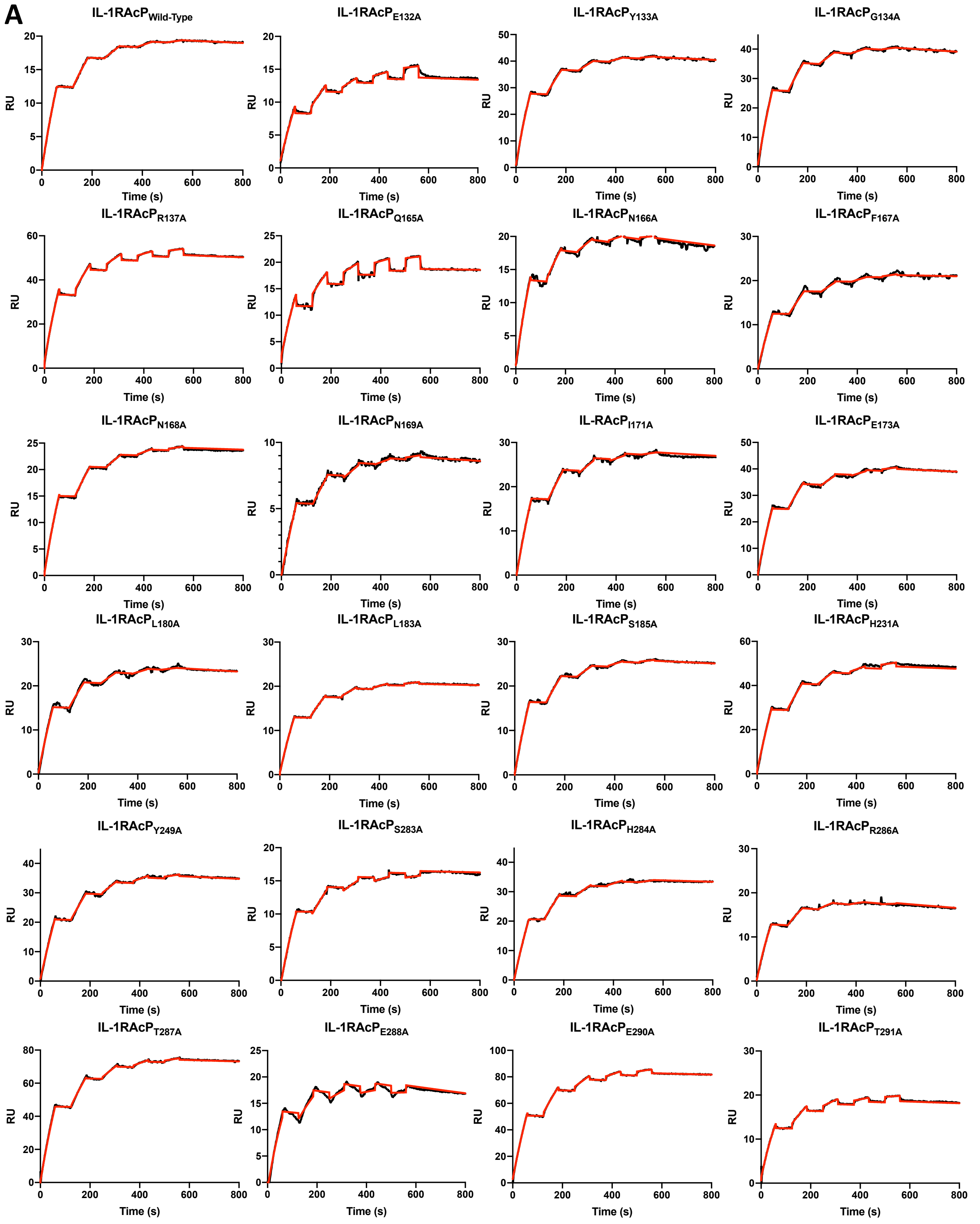

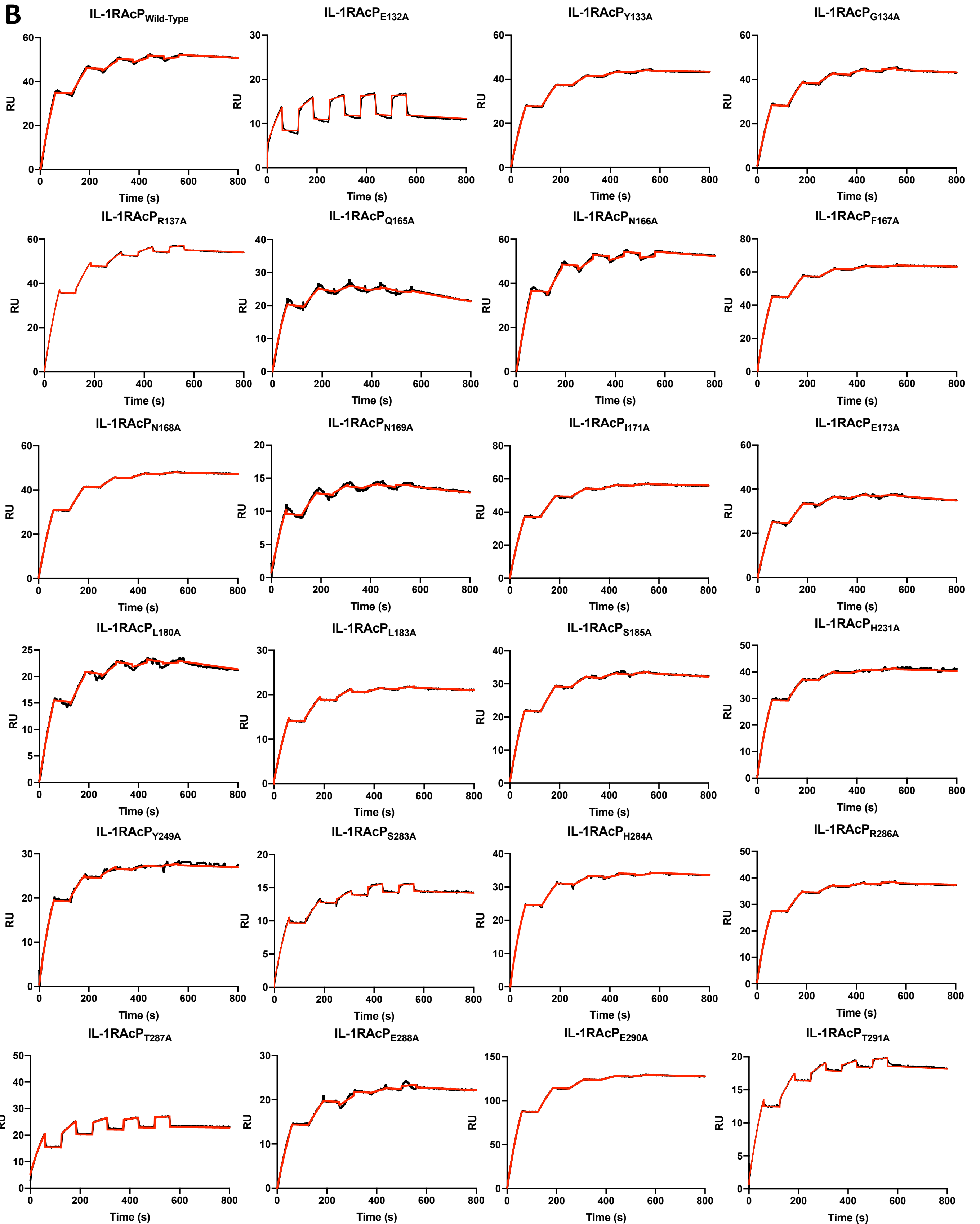
